# Suppression of aggregate and amyloid formation by a novel intrinsically disordered region in metazoan Hsp110 chaperones

**DOI:** 10.1101/2021.01.13.426581

**Authors:** Unekwu M. Yakubu, Kevin A. Morano

**Affiliations:** Department of Microbiology and Molecular Genetics, McGovern Medical School at UTHealth, Houston, TX USA; MD Anderson UTHealth Graduate School at UTHealth, Houston, TX USA

**Keywords:** Hsp70, Hsp110, chaperone, Alzheimer’s disease, Parkinson’s disease, amyloid, protein aggregation, proteostasis

## Abstract

Molecular chaperones maintain protein homeostasis (proteostasis) by ensuring the proper folding of polypeptides. Loss of proteostasis has been linked to the onset of numerous neurodegenerative disorders including Alzheimer’s, Parkinson’s, and Huntington’s disease. Hsp110 is related to the canonical Hsp70 class of protein folding molecular chaperones and interacts with Hsp70 as a nucleotide exchange factor (NEF), promoting rapid cycling of ADP for ATP. In addition to its NEF activity, Hsp110 possesses an Hsp70-like substrate binding domain (SBD) whose biological roles remain undefined. Previous work in *Drosophila melanogaster* has shown that loss of the sole Hsp110 gene (Hsc70cb) accelerates the aggregation of polyglutamine (polyQ)-expanded human Huntingtin, while its overexpression protects against polyQ-mediated neuronal cell death. We hypothesize that in addition to its role as an Hsp70 NEF, *Drosophila* Hsp110 may function in the fly as a protective protein “holdase”, preventing the aggregation of unfolded polypeptides via the SBD-β subdomain. Using an *in vitro* protein aggregation assay we demonstrate for the first time that *Drosophila* Hsp110 effectively prevents aggregation of the model substrate citrate synthase. We also report the discovery of a redundant and heretofore unknown potent holdase capacity in a 138 amino-acid region of Hsp110 carboxyl-terminal to both SBD-β and SBD-α (henceforth called the C-terminal extension). This sequence is highly conserved in metazoan Hsp110 genes, completely absent from fungal representatives, including *Saccharomyces cerevisiae SSE1*, and is computationally predicted to contain an intrinsically disordered region (IDR). We demonstrate that this IDR sequence within the human Hsp110s, Apg-1 and Hsp105α, inhibits the formation of amyloid Aβ-42 and α-synuclein fibrils *in vitro* but cannot mediate fibril disassembly. Together these findings demonstrate the existence of a second independent, passive holdase property of metazoan Hsp110 chaperones capable of suppressing both general protein aggregation and amyloidogenesis and raise the possibility of exploitation of this IDR for therapeutic benefit in combating neurodegenerative disease.

## Introduction

Molecular chaperones support cellular protein homeostasis (proteostasis) by assisting in the proper folding and assembly of nascent polypeptide chains, regulating activity and interactions of mature proteins, and shepherding damaged or short-lived proteins to degradation pathways (1). Environmental stressors such as oxidation, heat shock, or starvation are especially detrimental to incompletely folded proteins that have a propensity to misfold and form aggregates. In a balanced proteostatic network molecular chaperones such as Hsp70 and its co-factors prevent or reverse protein aggregation caused by partially or misfolded polypeptides (2, 3). However, with age and prolonged cellular damage, proteostasis becomes dysregulated leading to increased protein aggregation and resultant cytotoxicity (4–7).

Numerous neurodegenerative disorders such as Alzheimer’s, Parkinson’s, and Huntington’s disease have been linked to the misfolding and aggregation of key proteins in neuronal and muscle cells (4, 8). The misfolding of amyloidogenic polypeptides leads to the formation of structured oligomers that act as seeding elements for other amyloid-prone proteins as well as non-pathogenic polypeptides (9–11). Over time these amorphous oligomers form higher order energetically favorable and toxic amyloid fibrils which are difficult for the cell to disassemble. These occlusive fibrils, and likely the oligomeric intermediates that precede them, incorporate partially folded proteins, titrate chaperones away from nascent polypeptides, and contribute to overall cellular toxicity that ultimately results in neuronal cell death (5). Understanding the molecular mechanisms behind amyloidogenesis and its contribution to neurodegeneration can result in the development of therapeutics that halt or even reverse the progression of these devastating diseases. This is especially important given rapidly aging populations around the world, as many of these diseases, most notably Alzheimer’s disease, primarily affect persons aged 65 and older (5, 12).

The Hsp70 nucleotide exchange factor Hsp110 specifically plays a role in preventing the aggregation of neurodegenerative disease-associated polypeptides as demonstrated *in vitro* and *in vivo* with the use of animal models (13–16). The *Drosophila melanogaster* Hsp110 homolog Hsc70cb, when over-produced in fly lines also expressing the expanded poly-glutamine version of human Huntingtin (Htt) exon 1, prevents Htt aggregation and Huntington’s disease-associated phenotypes (17). Likewise, over-expression of Hsp110 mitigates protein aggregation and neurodegeneration in Parkinson’s and amyotrophic lateral sclerosis (ALS) mouse models (13, 18). Conversely, deletion of Hsp110 homologs in mice and *Drosophila* results in enhanced neurodegeneration resulting from increased amyloid formation of Alzheimer’s-linked Aβ42 and expanded polyglutamine protein, respectively (16, 19). Hsp110 is a member of the Hsp70 superfamily and as such shares the canonical Hsp70 architecture: an amino-terminal nucleotide binding domain followed by a short linker and a substrate binding domain (SBD) comprising two subdomains termed SBD-β and SBD-α (Figure 1A). SBD-β is a β-sandwich-containing putative substrate binding site, while SBD-α is an α-helical sequence that instead of clamping over the SBD-β region with associated peptide substrate as in all Hsp70 chaperones, binds to and stabilizes the Hsp110 NBD, promoting interaction with Hsp70 to form a stable heterodimer that facilitates nucleotide exchange (20–23).

**Figure 1:**
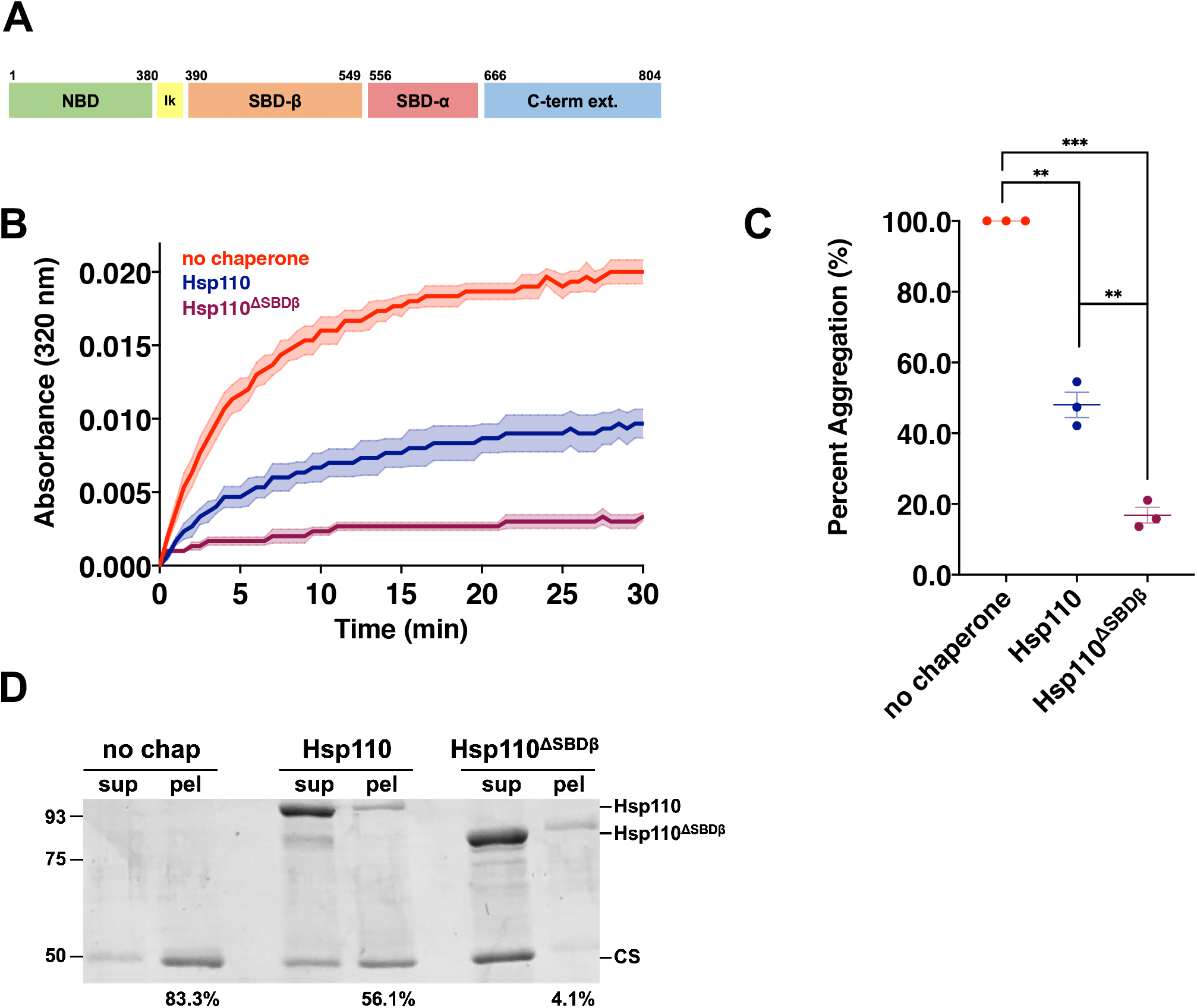
The *Drosophila* Hsp110 substrate binding domain, SBD-β, is expendable for aggregation suppression activity. **A**. Schematic of Hsp110 domain architecture: nucleotide binding domain (green) a.a. 1-380; linker (yellow) a.a. 381-389; substrate binding domain β (orange) a.a. 390-549; substrate binding domain α (red) a.a. 556-665; C-terminal extension (blue) a.a. 666-804 **B**. 200 nM denatured citrate synthase (CS) was incubated alone (no chaperone) or with 400 nM of respective chaperone: Hsp110, Hsp110^ΔSBDβ^ and light scattering measured at 320 nm, Bolded lines are the average of three replicates for each condition while the shaded region represents standard error of the mean (SEM). **C**. End point measurements of each condition were taken from Fig. 1B and divided by the no chaperone measurement within the respective replicate and converted to relative percentage. Group differences were analyzed using Welch’s *t*-test. *, p=0.05; **, p=0.005; ***, p=0.0005; ****, p=0.00005. **D**. Following light scatter assay endpoint samples were separated into soluble (sup) and insoluble (pel) fractions by differential centrifugation. Numbers indicate per cent of pellet signal, as quantified using ImageJ, from Coomassie-stained SDS-PAGE gel relative to combined sup plus pel signals.

While the NEF activity of Hsp110 has been well characterized, the biological role of substrate binding by the chaperone remains elusive (22, 24). *In vitro* studies have demonstrated that Hsp110 is a potent suppressor of protein aggregation in the absence of Hsp70, and SBD-β appears to dictate peptide binding in the yeast homolog Sse1 (25–28). Additionally, substrate binding by Sse1 is not required for the disaggregase activity exhibited by the Hsp40-Hsp70-Hsp110 triad (20). Moreover, Hsp110 appears to optimally function as an NEF at a 1:10 ratio with respect to Hsp70, significantly inhibiting Hsp70 activity at higher levels (14, 23, 29). This stoichiometry is inconsistent with the clear amyloid-suppressing effects observed upon overexpression in animal models of human neurodegenerative disease. Given the compelling protection from Htt-induced neurotoxicity provided by Hsp110 overexpression in flies we interrogated protein aggregation inhibition by fly Hsp110 through mutational analysis and revealed a previously undiscovered second substrate binding site in the extreme carboxyl-terminus. This region, henceforth referred to as the C-terminal extension, is absent in yeast Sse1 but conserved in Hsp110 genes from metazoans. We demonstrate that an intrinsically disordered region (IDR) within the C-terminal extension is both necessary and sufficient for classic passive chaperone holdase activity. Importantly, we show using thioflavin T binding assays and transmission electron microscopy that the IDR of both fly Hsp110 and human Hsp105α and Apg-1 Hsp110 homologs prevents the fibrilization of Aβ42 and α-synuclein, the highly amyloidogenic peptides involved in Alzheimer’s and Parkinson’s disease pathology, respectively. Our results support the conclusion that metazoan Hsp110 chaperones possess two distinct passive, yet potent, substrate binding activities and raise the possibility of their exploitation for therapeutic chaperone intervention in age-related neurodegenerative diseases.

## Results

### SBD-β is expendable for Hsp110 chaperoning activity

Previous studies by us and others have established the substrate binding capacity of the SBD-β subdomain *in vitro* using yeast and mammalian Hsp110 homologs without developing a clear understanding of the biological relevance of this chaperone property (26, 27, 30). To better explore the *in vivo* roles of Hsp110 substrate binding with respect to proteostasis and neurodegenerative disease we turned our attention to the fruit fly, *Drosophila melanogaster*, which possesses a single Hsp110 homolog encoded by the Hsc70cb gene (31–33). We purified the Hsc70cb-encoded protein with an added amino-terminal hexa-histidine tag (subsequently termed Hsp110) in *Escherichia coli* cells and utilized a standard spectrophotometric microplate light scattering assay to assess the ability of *Drosophila* Hsp110 to prevent protein aggregation of chemically denatured citrate synthase (CS) (Fig. S1A) (34). Hsp110 potently suppressed CS aggregation in a dose dependent manner with approximately 50% efficacy at a 1:1 ratio of chaperone to substrate. Stronger aggregate inhibition was observed at 2:1 and 4:1 ratios, indicating Hsp110 prevents aggregation in a manner similar to other homologs (Fig. S1B, S1C) (21, 27, 30, 35). We additionally validated the assay conditions by demonstrating that the non-specific protein bovine serum albumin (BSA) was unable to prevent CS aggregation (Fig. S2A). To verify that the increase in light scatter was indeed indicative of protein aggregation, we subjected samples obtained from the assay after 30 min to differential centrifugation to separate soluble and insoluble material. While both Hsp110 and BSA remained in the supernatant fraction, only Hsp110 was capable of preventing CS from entering the pellet (Fig. S2C).

To further understand the mechanism by which aggregation prevention occurs a series of deletion mutations were made in the SBD-β subdomain. Five deletion constructs were generated that removed portions or all of SBD-β, including the short hydrophobic linker connecting the nucleotide binding domain (NBD) to SBD-β (Fig. S3A). SBD-β deletion mutants were stably expressed, purified, and screened for chaperone activity (Fig. S3B-D). Surprisingly, we found that partial or complete removal of SBD-β had no significant deleterious effect on aggregation suppression by Hsp110 (Figs. 1B-C, S3C-D). Moreover, we noted that removal of SBD-β appeared to significantly enhance Hsp110 chaperoning activity to varying degrees. Interestingly, not only does removal of SBD-β fail to disrupt prevention of CS aggregation, but Hsp110^ΔSBDβ^ appears to be a much more potent holdase than full-length Hsp110 as demonstrated via differential centrifugation, with only 4% of citrate synthase remaining in the pellet fraction (Fig. 1D). To confirm that all our observations were not specific to the CS substrate used in the assays, we repeated the light scatter chaperone experiments using chemically-denatured firefly luciferase and obtained similar results, including enhanced chaperone activity at equivalent chaperone to substrate ratios (2:1) (Fig. S4). Together, these data indicate that the *Drosophila* Hsp110 SBD-β is expendable for holdase activity, suggesting the presence of an additional substrate binding site within the chaperone.

### The C-terminal extension of Hsp110 can operate independently of SBD-β to prevent protein aggregation through an intrinsically disordered region

Previous work using Chinese hamster (*Cricetulus griseus*) Hsp110 demonstrated that the NBD does not possess holdase activity as assessed using firefly luciferase (27). Consistently, we found that purified *Drosophila* Hsp110 NBD likewise was incapable of preventing aggregation of CS as demonstrated using both light scattering and differential centrifugation assays (Fig. 2). We noted that unlike the yeast Hsp110 Sse1, *Drosophila* Hsp110 possesses a carboxyl-terminal extension (C-term) of approximately 106 additional amino acids (Fig. S5). To assess whether this previously unstudied region may contribute to chaperone activity, we fused the C-term to the NBD and tested this purified protein for holdase function (Fig. S1A). Remarkably, the NBD-C-term hybrid protein exhibited potent aggregate suppression activity (Fig. 2A, B). Differential centrifugation confirmed the light scatter assay results, demonstrating that all of the NBD-C-term protein and the majority of unfolded CS remained in the soluble fraction after a 30 min incubation (Fig. 2C). Once again, we noted that like Hsp110^ΔSBDβ^, NBD-C-term appeared to be more efficient at aggregation prevention than full-length Hsp110 (Fig. 2). These findings demonstrate for the first time a second, previously unknown chaperoning site within a metazoan Hsp110 homolog that is capable of functioning independently of the canonical β-sandwich region (21, 27).

**Figure 2:**
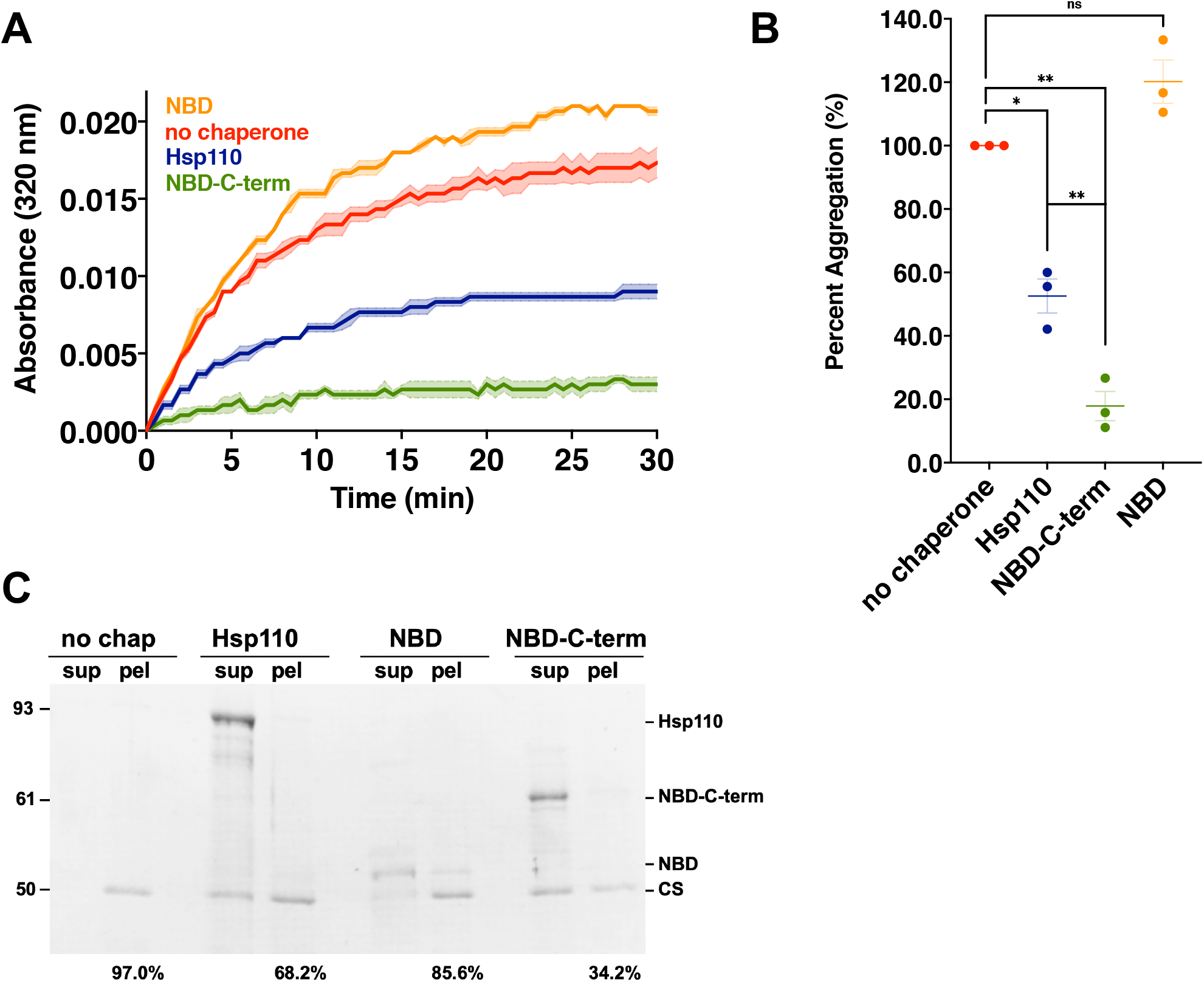
The far C-terminal extension of *Drosophila* Hsp110 is sufficient for aggregation suppression. **A**. 200 nM denatured CS was incubated alone (no chaperone) or with 400 nM of respective chaperone: Hsp110, NBD, NBD-C-term. Bolded lines are the average of three replicates for each condition while the shaded region represents standard error of the mean (SEM). **B**. End point measurements of each condition were taken from (Fig. 2A) and divided by the no chaperone measurement within the respective replicate and converted to relative percentage. Group differences were analyzed using Welch’s *t*-test. *, p=0.05; **, p=0.005; ***, p=0.0005; ****, p=0.00005. **C**. Following light scatter assay endpoint samples were separated into soluble (sup) and insoluble (pel) fractions by differential centrifugation. Numbers indicate per cent of pellet signal, as quantified using ImageJ, from Coomassie-stained SDS-PAGE gel relative to combined sup plus pel signals.

To more precisely map sequences conferring holdase activity within the C-term, we explored the *Drosophila* Hsp110 protein sequence to identify potential structural element boundaries to aid in design of additional deletions and truncations. As no full-length structures are available for *Drosophila* or any other metazoan Hsp110 homologs, we turned to *in silico* analysis. Using the protein disorder prediction package DISOPRED3, we identified two potential regions with high confidence scores for intrinsic disorder and the potential for protein binding (Fig. 3A) (36, 37). Of the two sites identified by the prediction software, one mapped to the extended loop region (residues 508-544) within SBD-β that differentiates this domain from the more distantly related Hsp70 superfamily homologs and another to the terminal 53 amino acids (residues 751-804) of the protein localized within the C-term region demonstrated to possess chaperone activity in Fig. 2. Previous studies have revealed that intrinsically disordered regions (IDRs) can act as sites of protein binding, assuming order when interacting with other proteins and peptides (38). This property is exploited by some protein chaperones. For example, the anti-aggregation/sequestration activity of the small heat shock protein (sHsp) family is mediated in part by IDRs (39–42). The localization of this IDR within the C-terminal extension made it a region of interest as a potential site for substrate-chaperone interaction. We therefore appended the predicted IDR to the NBD to create NBD-IDR and found that the IDR conferred robust aggregation prevention to the NBD (Fig. 3B, C). The holdase activity observed in the C-term domain of Hsp110 is thus due to the 53 amino acids composing a predicted IDR at the extreme carboxyl-terminus. Overall, these results confirm the presence of substrate binding capacity in the C-terminal extension mediated by a 53 amino acid intrinsically disordered region.

**Figure 3:**
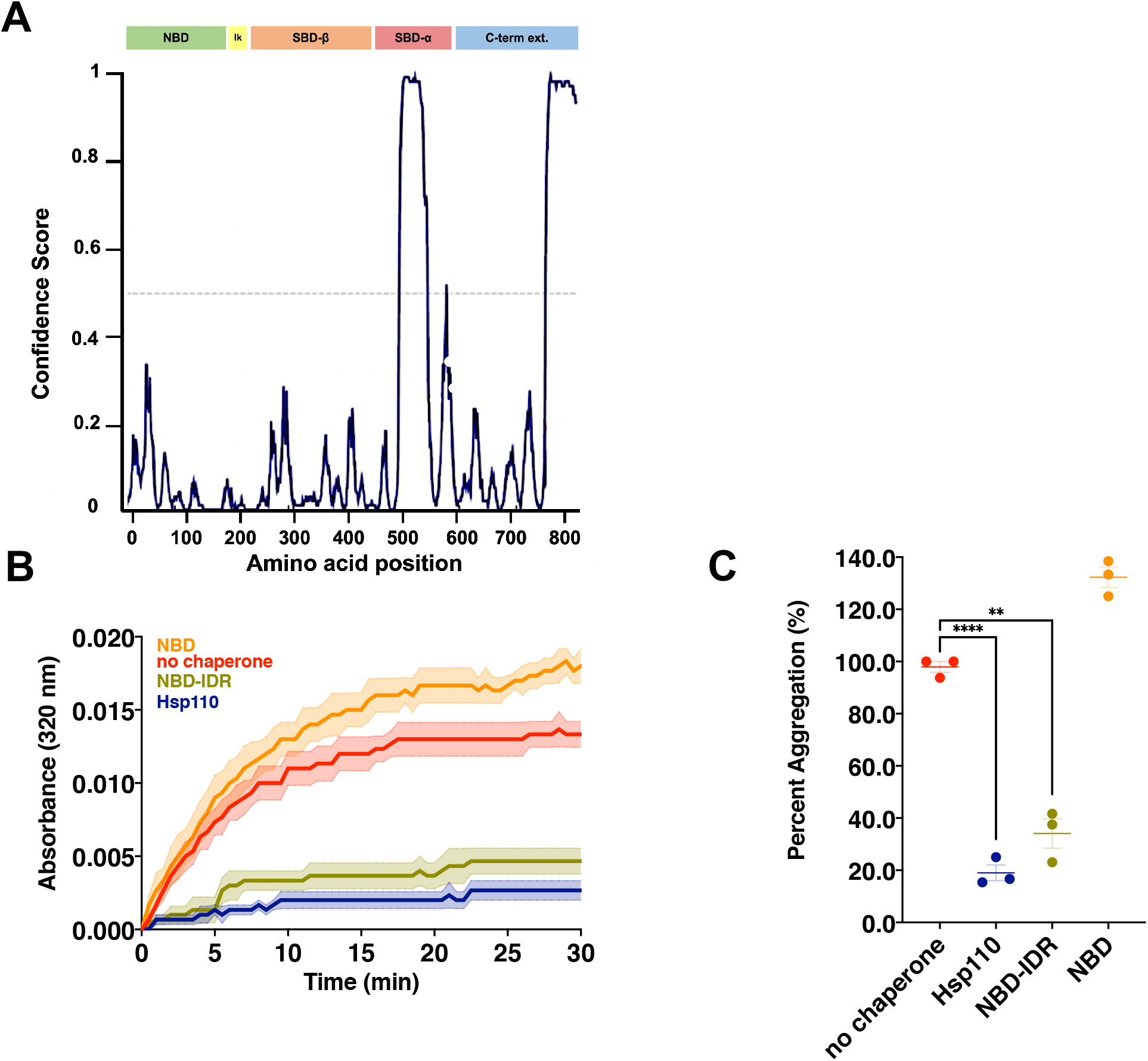
A predicted intrinsically disordered region (IDR) within the far C-terminal extension mediates aggregation suppression. **A**. Disorder prediction of *Drosophila* Hsp110 using DISOPRED3. Peaks above the dotted line (> 0.5 confidence score) indicate regions of predicted disorder. **B**. 200 nM denatured CS was incubated alone (no chaperone) or with 400 nM of respective chaperone: Hsp110, NBD, NBD-IDR. Bolded lines are the average of three replicates for each condition while the shaded region represents standard error of the mean (SEM). **C**. End point measurements of each condition were taken from (Fig. 3B) and divided by the no chaperone measurement within the respective replicate and converted to relative percentage. Group differences were analyzed using Welch’s *t*-test. *, p=0.05; **, p=0.005; ***, p=0.0005; ****, p=0.00005.

### The novel IDR-containing C-terminal extension is conserved in metazoan Hsp110s, Apg-1 and Hsp105α

Clustal Omega sequence alignment revealed that the C-terminal extension is present in several metazoan Hsp110 homologs including bovine, chimpanzee, orangutan, and the African clawed frog among others (Fig. S5) (43). Notably, the C-term extension and IDR appear to be conserved in the human Hsp110 homologs Apg-1and Hsp105α (Figs. S5 and 4A). While these regions are not strictly conserved among species in terms of sequence, there appears to be conservation in terms of amino acid characteristics and biochemical properties (44). Previous *in vitro* studies have demonstrated that both murine Hsp105α and Apg-1 possess passive chaperone holdases activity as we have discovered in the *Drosophila* Hsp110 (27, 45). We therefore constructed and purified chimeric versions of the NBD-C-term constructs in which the *Drosophila* Hsp110 NBD was fused to the C-terminal extensions of either human Hsp105α or Apg-1 (Fig. S1A). Using the light scatter CS aggregation assay, we found that both Apg-1 and Hsp105α C-terminal extensions, when fused to the *Drosophila* NBD, robustly prevented protein aggregation (Fig. 4B, C). Additionally, Apg-1 and Hsp105α C-terminal extensions appeared to be slightly more efficient holdases than full-length fly Hsp110. These results confirm the human Hsp110s contain conserved C-terminal extensions that act as potent passive chaperone regions, presumably through the IDR domains, and suggest potential evolutionary advantage of this region in animals.

**Figure 4:**
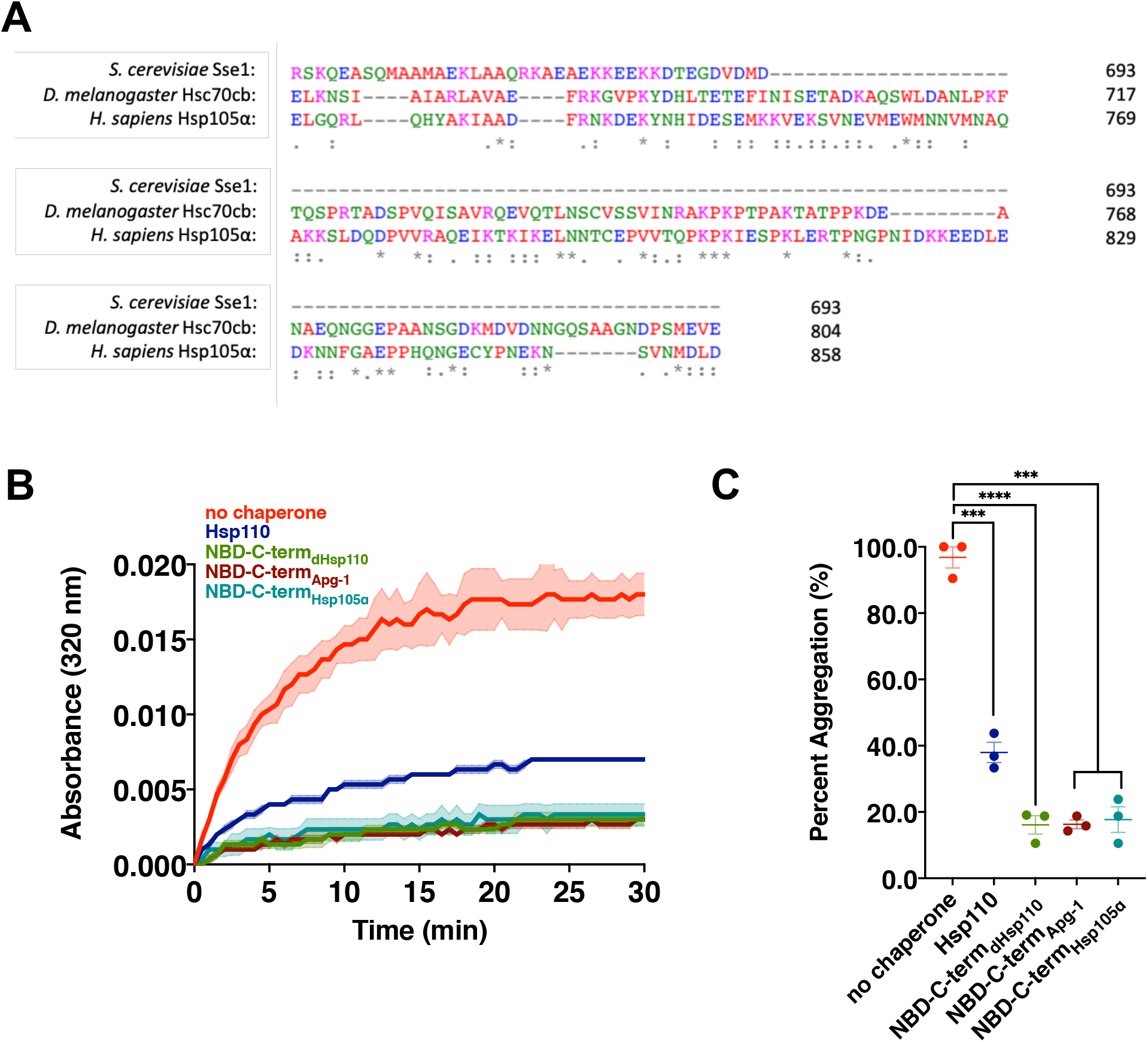
Aggregation suppression by conserved C-terminal extensions in human Hsp110 homologs Apg-1 and Hsp105α. **A**. Clustal Omega sequence alignment of the carboxyl-termini of Sse1 (budding yeast), Hsc70cB (fruit fly) and Hsp105α (human) Hsp110 homologs. **B**. 200 nM denatured CS was incubated alone (no chaperone) or with 400 nM of respective chaperone: Hsp110, NBD-C-term_dHsp110_, NBD-C-term_Apg-1_, NBD-C-term_Hsp105α_. Bolded lines are the average of three replicates for each condition while the shaded region represents standard error of the mean (SEM). **C**. End point measurements of each condition were taken from (Fig. 4B) and divided by the no chaperone measurement within the respective replicate and converted to relative percentage. Group differences were analyzed using Welch’s *t*-test. *, p=0.05; **, p=0.005; ***, p=0.0005; ****, p=0.00005.

### Human and *Drosophila* Hsp110 IDRs prevent amyloid formation

The presence of the C-terminal extension and the conservation of this substrate binding capacity in human Hsp110s prompted us to ask what potential it has for acting on proteins implicated in human neurodegenerative disease linked to protein conformation (13, 16–18, 46). Amyloid β (Aβ) is a cleavage product of the amyloid precursor protein (APP), and a major component of amyloid plaques associated with Alzheimer’s disease toxicity (47–49). APP cleavage creates two forms of amyloid β peptide, Aβ42 and Aβ40, of which Aβ42 is the more toxic and amyloidogenic form (50). Aβ peptides will assemble over time into oligomeric species that ultimately form large fibrils that comprise the amyloid plaques (51, 52). The fibrilization reaction can be followed over time *in vitro* by assessing binding of the fluorescent dye thioflavin T, which selectively binds β-sheets in amyloid structures (53, 54). We therefore developed a microplate thioflavin T-based fibrilization assay and observed significant signal production from commercially obtained Aβ42 monomers over a 16 hr time course (Fig. 5A, B). Addition of the NBD had little impact on fibrilization yield or kinetics. We generated two new chimeric protein fusions in which only the predicted IDR of both *Drosophila* Hsp110 and human Hsp105α were fused to the *Drosophila* NBD and found that in contrast to NBD alone, both the IDR fusions completely prevented the fibrilization of Aβ42. Consistent with previous reports, the addition of a Hsp110 chaperone, in this case the minimal chaperoning entity NBD-IDR_Hsp105α_, in the absence of Hsp70 and Hsp40 did not resolubilize fibrils that have already formed (Fig. S6) (14, 45).

**Figure 5:**
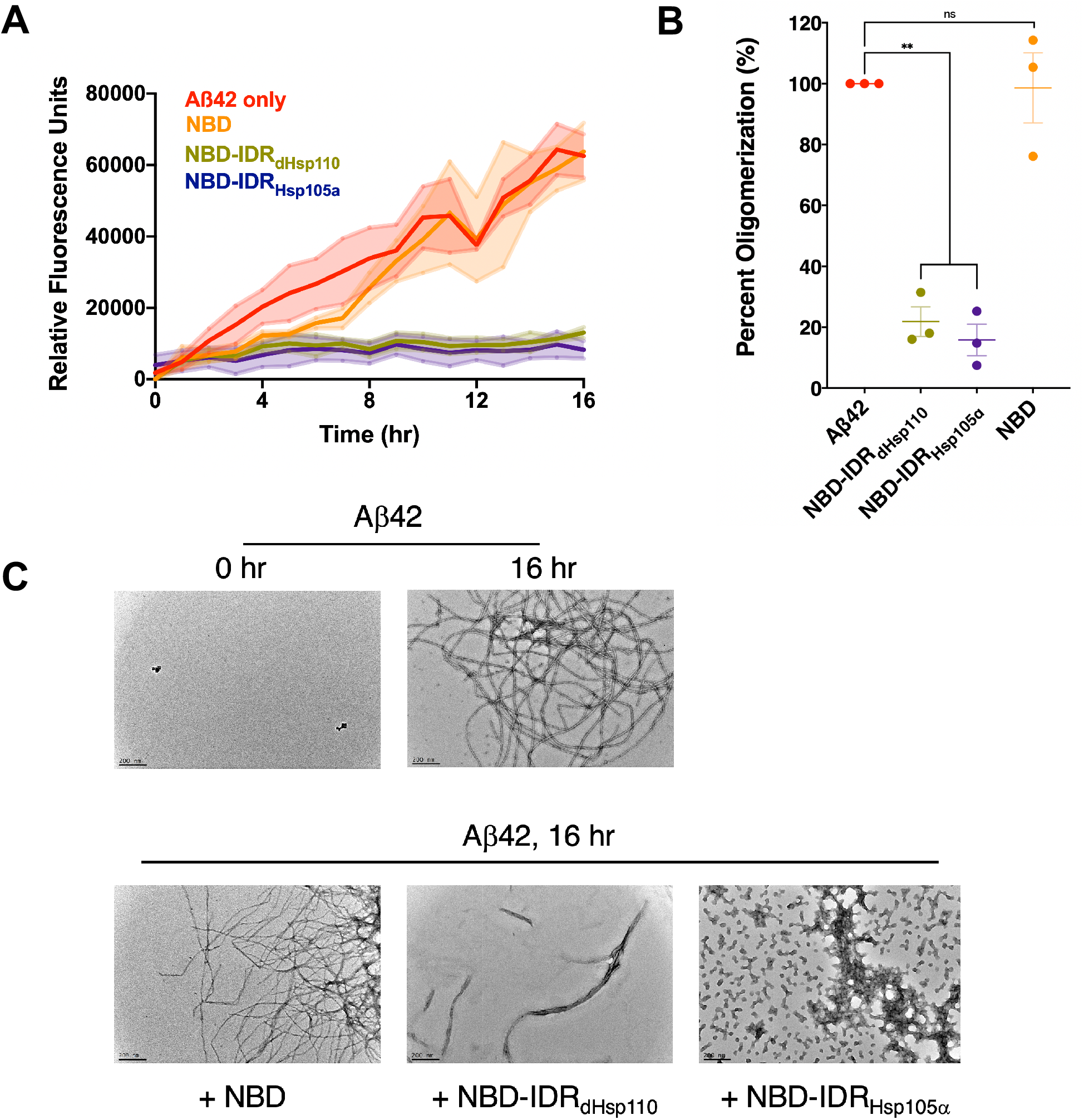
*Drosophila* and human IDRs prevent oligomerization and fibril formation of Alzheimer’s peptide amyloid β. **A**. 2 μM of Aβ42 was incubated alone (Aβ42 only) or with 4 μM of respective chaperone: NBD, NBD-IDR_dHsp110_, NBD-IDR_Hsp105α_ in the presence of the amyloid-detecting dye thioflavin T for the indicated time course and fluorescence detected as described in Materials and Methods. Bolded lines are the average of three replicates for each condition while the shaded region represents standard error of the mean (SEM). **B**. End point measurements of each condition were taken from (Fig. 5A) and divided by the no chaperone measurement within the respective replicate and converted to relative percentage. Group differences were analyzed using Welch’s *t*-test. *, p=0.05; **, p=0.005; ***, p=0.0005; ****, p=0.00005. **C**. Endpoint samples from the thioflavin T binding assay were recovered, negative stained and imaged via transmission electron microscopy, with or without additional chaperones as indicated. Scale bar= 200 nm.

Aggregation and plaque formation of the protein α-synuclein (α-syn) is a major contributor to Parkinson’s disease. Fibrilization of α-syn can also be tracked via thioflavin T fluorescence but typically requires seeding by preformed fibrils (PFF) in *in vitro* assay systems. We obtained commercially prepared α-syn monomers and PFF and observed modest thioflavin T fluorescence over a 24 hr time course from the PFF but not the monomers (Fig. S7). Addition of PFF to α-syn monomers resulted in a slow increase in thioflavin T fluorescence over 24 hr in a manner that was accelerated by the further addition of the *Drosophila* NBD by an unknown mechanism with no difference in overall signal yield. Remarkably, both NBD-IDR fusions allowed a small increase in fluorescence over the first hour of the experiment but little further increase, demonstrating highly effective suppression of α-syn fibrilization (Fig. S7B, C).

To further assess the ability of C-terminal IDRs to prevent fibrilization transmission we turned to transmission electron microscopy (TEM), which allows visualization of amyloid Aβ42 structures. Samples were removed from the microplate wells after the 16 hr thioflavin T experiment and deposited onto grids for TEM. Extensive fibrilization of Aβ42 was observed after 16 hr that was absent at the initiation of the assay (Fig. 5C, top panels). Incubation of Aβ42 with both NBD-IDRs drastically decreased formation of higher order Aβ42 fibrils while addition of the NBD alone had no impact on fibril formation (Fig. 5C, bottom panels). The fibrils that did form in the presence of the IDRs were predominantly shorter and smaller than the fibrils seen with Aβ42 at 16 hr. We noted that in the presence of NBD-IDR_Hsp105α_, but not NBD-IDR_dHsp110_, numerous small globular structures were observed. To further investigate this phenomenon, we incubated NBD-IDR_Hsp105α_ alone for 16 hr under the same conditions and observed the same particles which averaged approximately 30 nm in diameter, indicating that the chaperone fusion alone is capable of forming these assemblies (Fig. S8). Together these experiments establish that the C-terminal IDRs of *Drosophila* and human Hsp110 homologs are capable of inhibiting fibrilization of two critical amyloidogenic substrates *in vitro*, strongly suggesting that this domain may impart similar activity to the Hsp110 in flies and humans.

## Discussion

In this study we reveal a novel aggregation prevention mechanism in metazoans provided by a previously unstudied region at the far C-terminus of the Hsp110 family of molecular chaperones. This anti-aggregation property is mediated by predicted intrinsically disordered regions within the C-terminal extension and includes amino acids 718-804 in *Drosophila* Hsc70cb and 803-858 in human Hsp105α. Additionally, IDR-mediated chaperoning can prevent amyloid formation of Aβ42 and α-synuclein, the causative agents of Alzheimer’s and Parkinson’s diseases, respectively. These functions do not require the presence of the canonical SBD-β substrate binding domain, establishing the presence of a second chaperone site within this member of the Hsp70 chaperone superfamily.

Intrinsically disordered regions are protein segments that do not maintain a fixed structure in their native states (55). IDRs tend to be low in amino acid complexity and sequence conservation. Nonetheless, their abundance in eukaryotic species indicates they serve an essential purpose (56). IDRs have been shown to assume secondary and tertiary structure when interacting with other polypeptides (57). The abundance of IDRs in nature, particularly in higher order organisms, has prompted investigation into the functional relevance of these sites in protein-protein interactions (58). Previous work supports the function of small heat shock proteins (sHsps) being due in part to IDRs facilitating chaperoning activities (38, 59). sHsps including yeast Hsp42 act as sequestrases, corralling misfolded proteins to prevent their aggregation until the Hsp70 machinery can bind and facilitate proper folding conformations or target substrates to degradation pathways (60). In a similar manner, the Hsp110 NBD-IDR constructs appear to function by binding to unfolded and amyloidogenic proteins in an ATP-independent manner, preventing aggregation and maintaining these proteins in a soluble state. It is possible the IDR in full-length Hsp110 provides a similar function in addition to the well-established nucleotide exchange activity characteristic of this family of chaperones.

While the resolution of the light scatter assays is somewhat limited, it appears that chimeric proteins consisting of either the entire C-terminal extension or just the IDR appended to non-chaperoning NBD prevent aggregation in a capacity similar to, and in some cases better than, full-length Hsp110. It is unclear why this is the case, but we speculate it may be due to differential accessibility of the IDRs. It is possible the folding of full-length Hsp110 is such that it partially obscures the C-terminal extension, making it harder to fully interact with misfolded polypeptides. Interestingly, removal of some or all of SBD-β also enhanced inhibition of CS aggregation in the context of the otherwise full-length Hsp110 protein (Fig. 1B, S3C). Deletion of this domain may cause changes in protein structure such that the IDR is fully exposed and more available to interact with substrates. Alternatively, SBD-β itself may interact with the IDR or otherwise occlude the IDR from productive substrate interaction. An interesting ancillary observation is the assembly of NBD-IDR_Hsp105α_ into what appear to be globular particles. Initially we speculated that these images represented NBD-IDR_Hsp105α_ oligomerizing in concert with Aβ42 oligomers or protofibrils. However, NBD-IDR_Hsp105α_ clearly assembled into the same particles in the absence of Aβ42 (Fig. S8). Additionally, Aβ42 is typically found in fibril rather than spherical oligomeric states, making it unlikely that the substrate is dictating the NBD-IDR ultrastructure. TEM images of the classic sHsp αB-crystallin highly resemble the NBD-DR particles, with similar structure and diameter (61). It is therefore possible that these particles represent an alternative native conformation of NBD-IDR_Hsp105α_ that is not a prerequisite for chaperone activity as we did not observe the same behavior for NBD-IDR_dHsp110_. As there are no reports of native human Hsp110 chaperones assembling into such particles, it is also likely that they are a byproduct of an “unrestricted” IDR.

We have demonstrated that the C-terminal IDRs are potent suppressors of Aβ42 and α-synuclein oligomerization *in vitro*. Overexpression of Hsp110 in several different biological systems has been reported to reduce formation of toxic misfolded protein conformers (13, 15– 18). These findings are at odds with the known inhibitory effect that Hsp110 has on Hsp70 ATPase activity and thus protein folding when present above a roughly 1:10 ratio (110:70) due to excessive NEF activity. Moreover, our *in vitro* experiments firmly establish that the Hsp110 C-terminal IDR is capable of suppressing aggregate and amyloid formation in the absence of Hsp40 and Hsp70. Thus, while modulation of Hsp70 substrate cycling may be one mechanism by which Hsp110 contributes to protein homeostasis, another Hsp70-independent mechanism clearly exists. We are currenting attempting to build *in vivo* evidence for roles for both SBD-β and the C-terminal IDR in metazoan neurogenerative disease onset and progression using *Drosophila* amyloidogenesis models.

It is tempting to speculate that our findings suggest a potential functional therapeutic for human neurodegenerative disease. Engineered chaperone overexpression to combat diseases of protein misfolding is fraught with non-specific and collateral effects due to the key roles chaperones play in all cellular biology (62–64). It is possible that further investigations into using the Hsp110 IDR peptide alone or attached to an inert protein scaffold may yield a deliverable tool that would block or delay fibril and plaque formation in several amyloidogenic disorders.

## Materials and Methods

### Plasmid construction

Hsc70cb cDNA was amplified using PCR and inserted into the pProEX-HTA protein expression vector using 5’ SacI and 3’ SpeI restriction enzyme sites (Invitrogen, Carlsbad, CA, USA). Two *Drosophila* NBD constructs (with or without a stop codon) were created by PCR amplification followed by cloning into pProEX-HTA using 5’ SacI and 3’ SpeI restriction sites. The *Drosophila* NBD-C-term construct was designed and ordered from Genewiz (South Plainfield, NJ) and subcloned into pProEX-HTA using 5’ SacI and 3’ SpeI restriction sites. Apg-1 and Hsp105α NBD-C-term constructs were created by PCR amplification of the respective C-terminal extensions from cDNA and insertion into the nonstop *Drosophila* pProEX-HTA-NBD plasmid using 5’ Spe1 and 3’ XhoI restriction sites. NBD-IDR constructs were created by PCR amplification of Hsc70cb and Hsp105α C-terminal IDRs, followed by cloning into the non-stop *Drosophila* pProEX-HTA-NBD plasmid using 5’ SpeI and 3’ XhoI restriction sites.

### Protein purification

BL21 *Escherichia coli* cells were used to express and purify all chaperones and fusion proteins used in this study. Sub-culture of overnight inoculum was made to an OD of 0.15 in 600 mL Luria Broth with ampicillin (100 mg/mL) and grown to OD 0.6. Isopropyl β-d-1-thiogalactopyranoside was added to the culture flask to a final concentration of 1 mM and flasks were subject to shaking with aeration for 4 hr at 25°C. Cell pellets were collected and flash frozen until processing for immobilized affinity chromatography (iMAC). For iMAC, cells were thawed and lysed using chemical lysis Buffer B (50 mM Tris Base - pH 7.5, 5 mM imidazole, 2 mM MgCl_2_, 200 mM NaCl, 10% octylthioglucoside). Lysates were incubated with Talon cobalt resin (TakaraBio, USA, Mountain View, CA) for 1 hr at 4°C with top-over-bottom mixing. Following incubation, resin was separated from lysates using centrifugation, washed with Buffer B and Buffer C (50 mM Tris - pH 7.5, 10 mM imidazole, 2 mM MgCl_2_, 600 mM NaCl), and chaperones eluted with Buffer E (50 mM Tris - pH 7.5, 200 mM imidazole, 2 mM MgCl_2_, 700 mM NaCl). Eluates were concentrated to ∼500 μL and kept at −80°C until further purification via size exclusion chromatography (SEC). iMAC samples were thawed on ice and loaded into a purification column packed with 10 mL of Sephacryl S-100 (GE Healthcare) equilibrated with 25 mM Tris-HCl, pH 7.5 100 mM NaCl buffer. Samples were eluted in the same buffer and peak fractions collected and frozen at −80°C until use.

### Light scatter aggregation assay

Stock citrate synthase (Sigma-Aldrich, St. Louis, MO) and firefly luciferase solutions were diluted to 18.7 μM using a chemical denaturing buffer (6.6 M guanidine hydrochloride, 5.5 mM dithiothreitol) and incubated for 1 hr at 25°C before assays. Aggregation assays were performed by incubating 200 nM denatured citrate synthase or firefly luciferase with 400 nM of respective chaperones, chimeric proteins or bovine serum albumin in a refolding buffer (25 mM Tris-HCl, pH 7.5, 100 mM NaCl) in a total volume of 180 μL in a clear-bottom 96-well plate. An MX Synergy (BioTek, Winooski, VT) plate reader was used to obtain 320 nm absorbance readings every 30 sec for 30 min (CS), or 90 minutes for firefly luciferase, at 25°C. Assays were performed in technical triplicate for each experiment and averaged for three separate experimental replicates.

### Differential centrifugation

To quantify the amount of soluble versus insoluble protein after the aggregation assays, 175 μL of the reaction was taken at the conclusion of the time course. This fraction was subjected to differential centrifugation at 16,000 x*g* for 4 min. The supernatant (169 μL) was removed to a new tube and the remaining pellet fraction was brought to 169 μL with refolding buffer. Both supernatant and pellet fractions were brought to 400 μL volume with molecular bology-grade water and 40 μL of trichloroacetic acid was added to precipitate proteins. After a 45 min incubation samples were spun at 16,000 x*g* for 15 min, washed with acetone, dried and resolubilized in 15 μl SDS-PAGE sample buffer and fractions analyzed using SDS-PAGE and Coomassie staining. Amount of substrate in each fraction as a percent of total was quantified using ImageJ software (NIH).

### Thioflavin T binding assay

α-synuclein thioflavin T binding assays were performed by incubating 2 μM α-synuclein monomer (StressMarq Biosciences, Victoria, BC, Canada), 1 μM pre-formed fibrils (StressMarq Biosciences, Victoria, BC, Canada), 4 μM of each respective chaperone, and 5 μM thioflavin T (Sigma, St. Louis, MO) in a total reaction volume of 180 μL using a 96-well μClear bottom black plate (Greiner Bio-One, Monroe, NC). Samples were subject to excitation at 450 nm and emission at 490 nm at 37°C and fluorescence intensity was measured every five min for 24 hr using an MX Synergy microplate reader. Aβ42 was purchased from GenScript USA (Piscataway, NJ) and thioflavin T binding assays were performed by incubating 2 μM Aβ42, 4 μM of each respective chaperone, and 5 μM thioflavin T in a total reaction volume of 50 μL in a 96-well μClear bottom black plate. Fluorescence intensity measurements (ex. 450/em. 490) were taken every 5 min for 16 hr at 37°C using an MX Synergy microplate reader.

### Disaggregation assay

Aβ42 disaggregation assay was performed by incubating 2 μM Aβ42 and 5 μM thioflavin T in refolding buffer in a total reaction volume of 50 μL in a 96-well μClear bottom plate. Fluorescence intensity measurements (ex. 450/ em. 490) were taken every 5 minutes for 20 hr at 37 °C using a BioTek MX Synergy microplate reader. At 20 hr, 4 μM of NBD-IDR_Hsp105α_ was added and the reaction continued for an additional 6 hr.

### Transmission electron microscopy

Samples were prepared by taking baseline fluorescence measurements of all conditions and removing Aβ_1-42_ at T=0. The remaining samples were measured by the protocol detailed in the thioflavin T binding assay section after which samples were recovered and diluted to a final protein concentration of 1.0 μM. Samples were negatively stained with 1% uranyl acetate in Tris NaCl refolding buffer and fixed onto 400 mesh copper grids (Electron Microscopy Sciences, Hatfield, PA). TEM imaging was done at 100x magnification with a JEOL 1400 electron microscope. Representative images were used for all conditions. Measurement of particles observed with the NBD-IDR_Hsp105α_ chimeric protein was performed using the distance tool on pixel-calibrated images using ImageJ.

## Statistical analysis

Welch’s variance *t*-test of unequal variance was used to analyze the mean differences between conditions. End point measurements for each replicate in a condition were averaged and analyzed using Prism 9 (Graphpad Software, San Diego, CA). For all time course experiments measurements at each time point were taken and averaged and the standard error of the mean was calculated. For the plots darker solid lines represent the calculated mean while the light shading represents the standard error of mean for those data. For all significance tests, *, p=0.05; **, p=0.005; ***, p=0.0005; ****, p=0.00005.

## Acknowledgements

We thank Drs. Bo Hu, Pratick Khara and Kohei Kishida (Department of Microbiology and Molecular Genetics, McGovern Medical School) for expert assistance with transmission electron microscopy image acquisition and interpretation. We thank Dr. Sheng Zhang from the McGovern Medical School Institute of Molecular Medicine for providing the Hsc70cb expression plasmid. This study was supported by the National Institutes of Health (GM127287) and the Russell and Diana Hawkins Family Foundation Discovery Fellowship from the MD Anderson UTHealth Graduate School of Biomedical Sciences.

## Author contributions

UMY and KAM planned and outlined the scope of the work and manuscript. UMY wrote the initial drafts and was responsible for all revisions. KAM edited the manuscript. UMY and KAM designed and created the figures.

---

**Figure S1:**
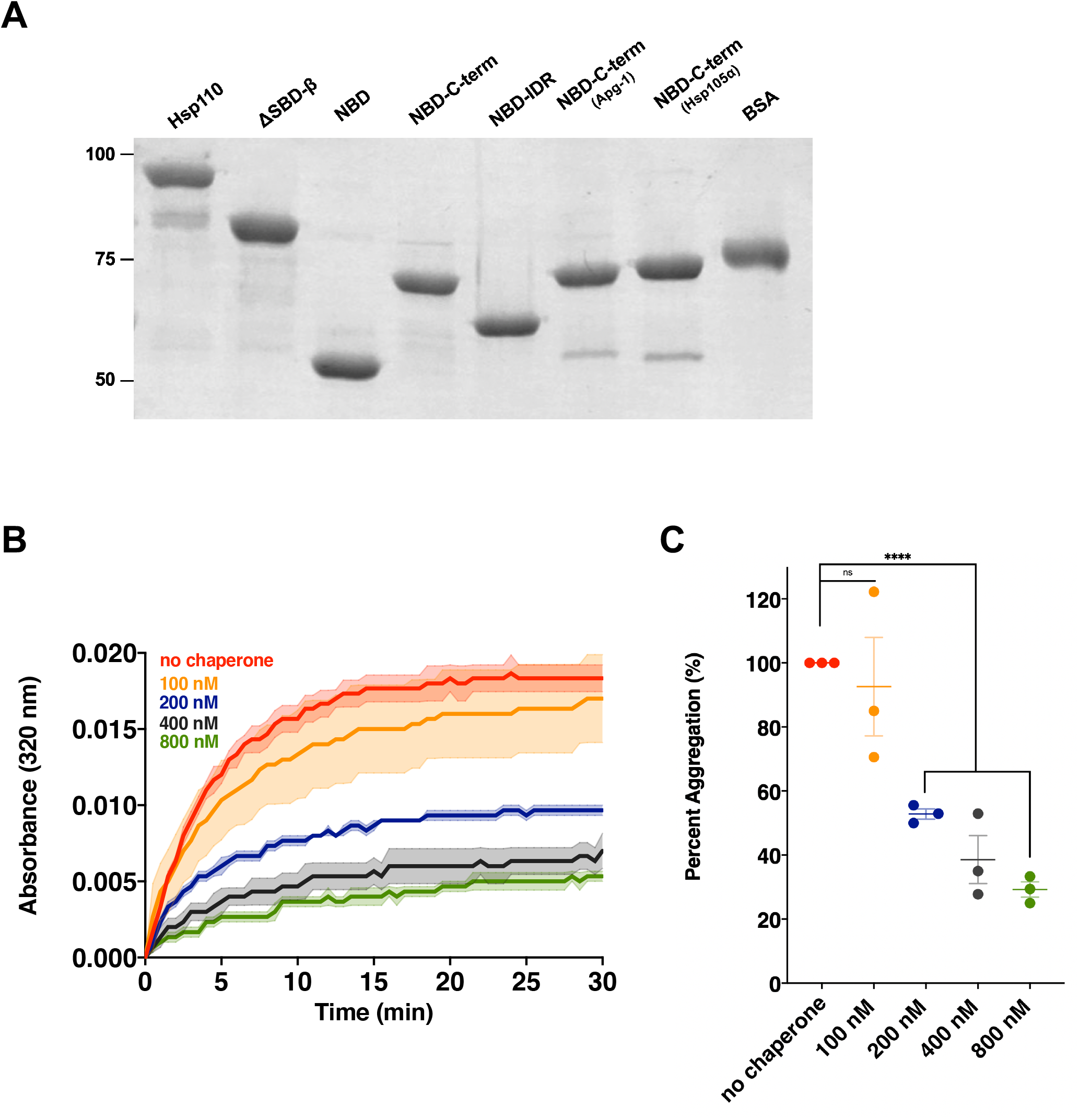
Hsp110 prevents aggregation in a dose-dependent manner. **A**. 500 ng of each purified chaperone variant used in this study shown via 12% SDS-PAGE stained with Coomassie Brilliant Blue. **B**. 200 nM denatured CS was incubated alone (no chaperone) or with *Drosophila* Hsp110 at .**5x, 1x, 2x**, or **4x** concentration. Bolded lines are the average of three replicates for each condition while the shaded region represents standard error of the mean (SEM). End point measurements of each condition were taken from (Fig. S1B) and divided by the no chaperone measurement within the respective replicate and converted to relative percentage. Group differences were analyzed using Welch’s *t*-test. *, p=0.05; **, p=0.005; ***, p=0.0005; ****, p=0.00005.

**Figure S2:**
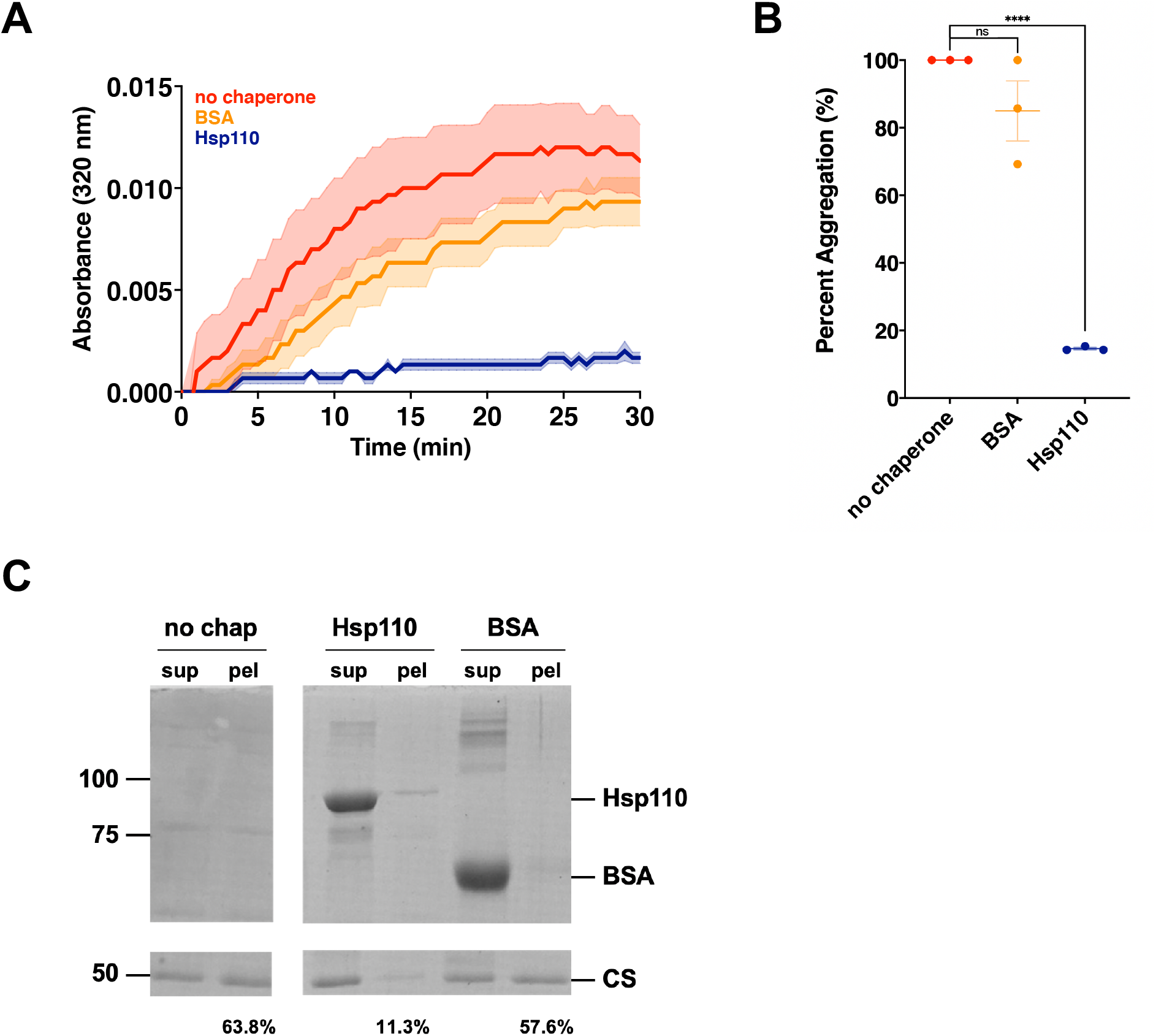
Non-chaperone protein bovine serum albumin (BSA) does not prevent CS aggregation. **A**. 200 nM denatured CS was incubated alone (no chaperone) or with 400 nM Hsp110 or BSA. **B**. End point measurements of each condition were taken from (Fig. S2A) and divided by the no chaperone measurement within the respective replicate and converted to relative percentage. Group differences were analyzed using Welch’s *t*-test. *, p=0.05; **, p=0.005; ***, p=0.0005; ****, p=0.00005. **C**. Following light scatter assay endpoint samples were separated into soluble (sup) and insoluble (pel) fractions by differential centrifugation. Numbers indicate per cent of pellet signal, as quantified using ImageJ, from Coomassie-stained SDS-PAGE gel relative to combined sup plus pel signals.

**Figure S3:**
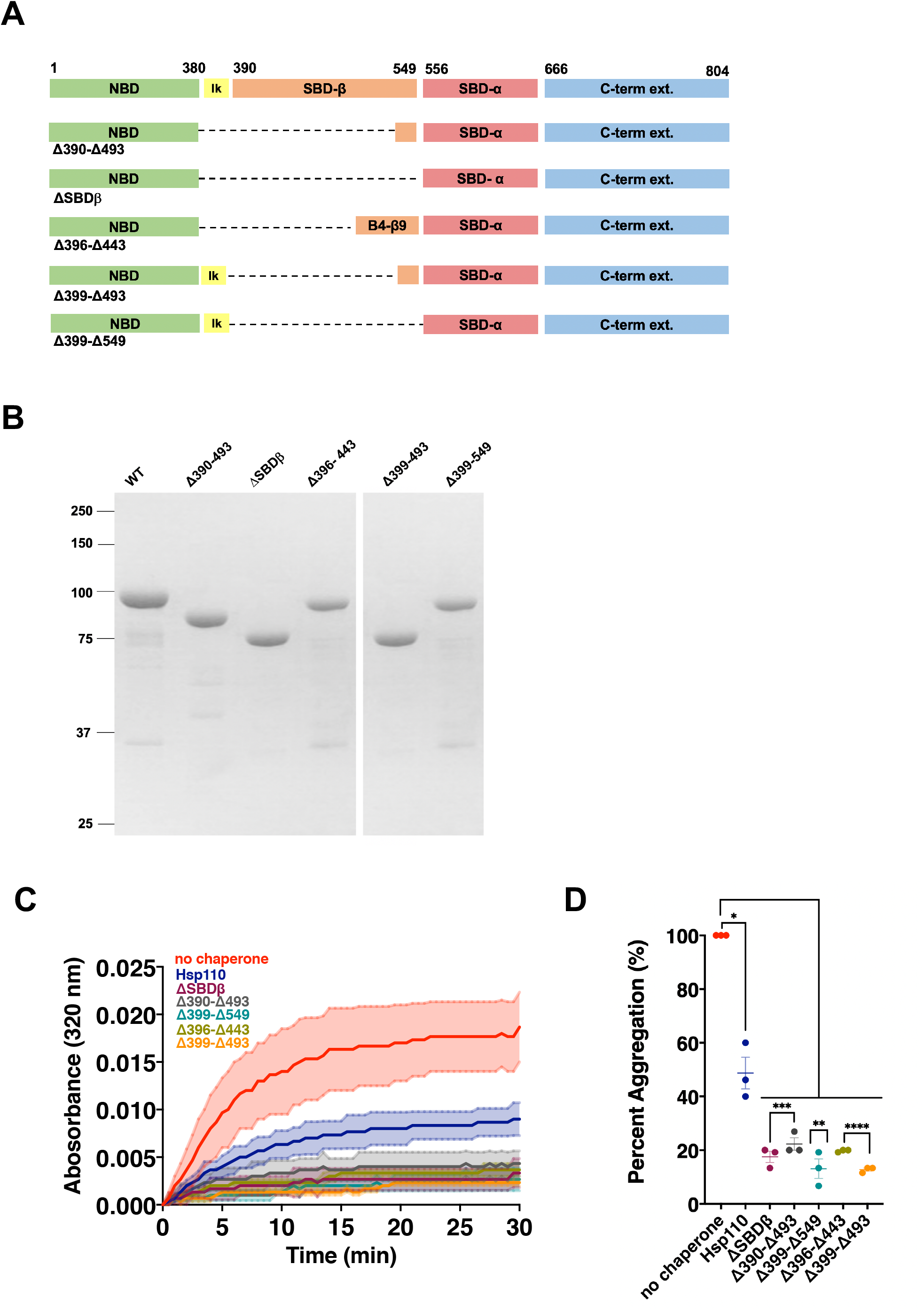
Partial or complete deletion of SBD-β does not eliminate Hsp110 substrate binding activity. **A**. Schematic of Hsp110 domain architecture indicating various SBD-β deletion mutants. **B**. 200 ng of purified Hsp110 and each SBD-β deletion mutant used in this study via 12% SDS-PAGE stained with Coomassie Brilliant Blue. **C**. 200 nM denatured CS was incubated alone (no chaperone) or with 400 nM of respective chaperone: Hsp110, Hsp110^ΔSBDβ^, Hsp110^Δ390-493^, Hsp110^Δ399-549^, Hsp110^Δ396-443^, Hsp110^Δ399-493^. Bolded lines are the average of three replicates for each condition while the shaded region represents standard error of the mean (SEM). **D**. End point measurements of each condition were taken from (Fig. S3C) and divided by the no chaperone measurement within the respective replicate and converted to relative percentage. Group differences were analyzed using Welch’s *t*-test. *, p=0.05; **, p=0.005; ***, p=0.0005; ****, p=0.00005.

**Figure S4:**
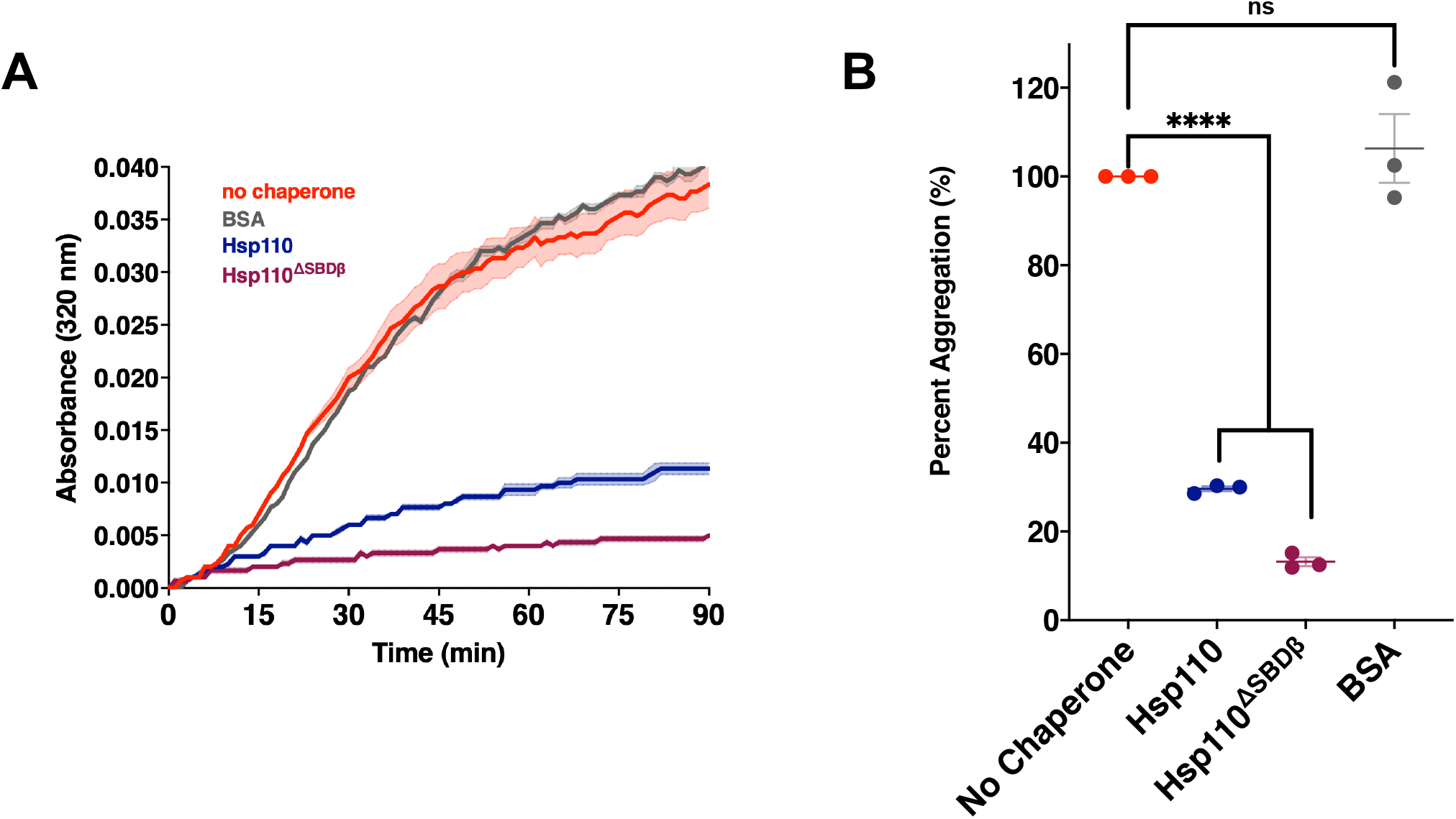
SBDβ is not required for aggregation prevention of denatured firefly luciferase. **A**. 200 nM denatured firefly luciferase was incubated alone (no chaperone) or with 400 nM of respective chaperone/protein: Hsp110, Hsp110^ΔSBDβ^, BSA. Bolded lines are the average of three replicates for each condition while the shaded region represents standard error of the mean (SEM). **B**. End point measurements of each condition were taken from (Fig. S4A) and divided by the no chaperone measurement within the respective replicate and converted to relative percentage. Group differences were analyzed using Welch’s *t*-test. *, p=0.05; **, p=0.005; ***, p=0.0005; ****, p=0.00005.

**Figure S5:**
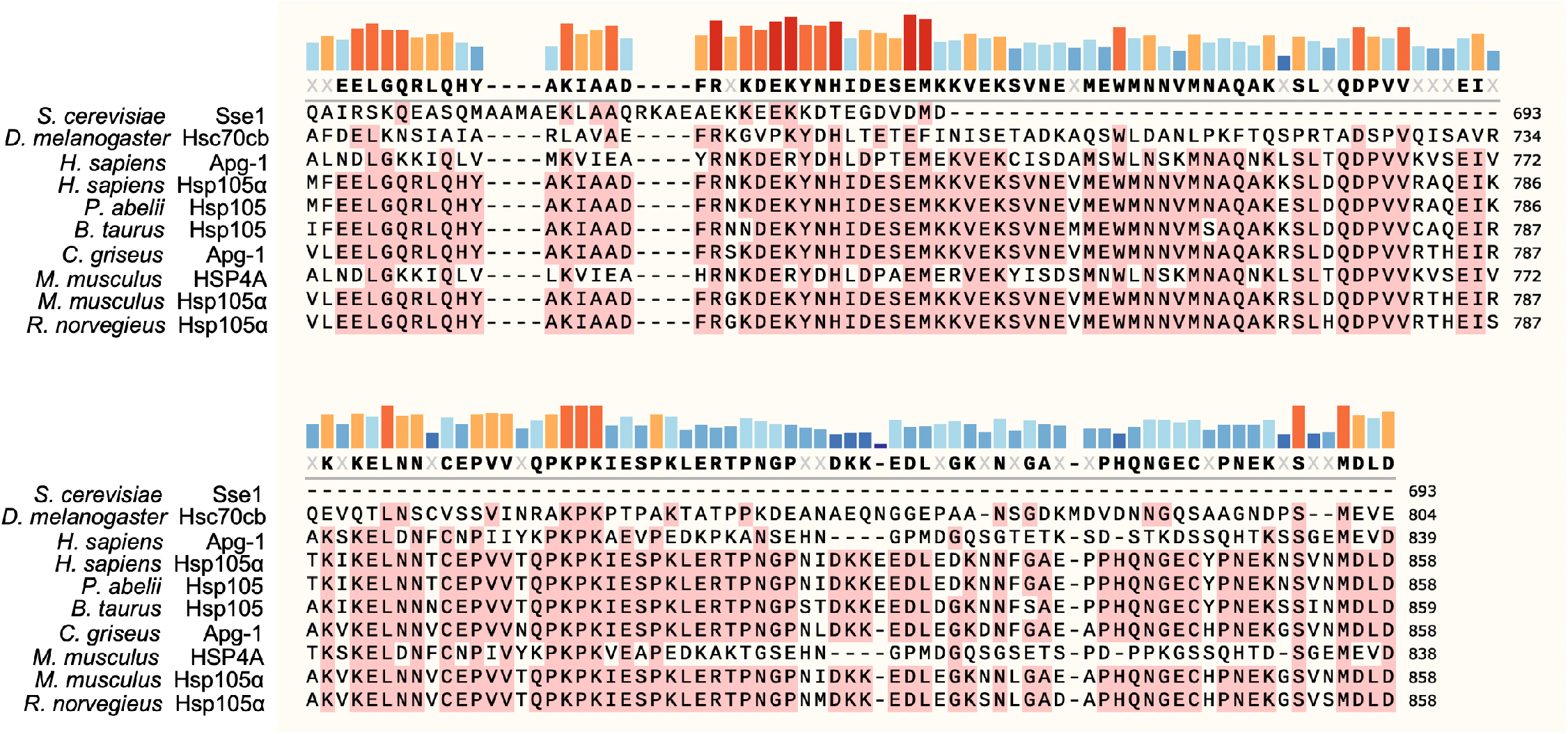
Multiple species alignment of Hsp110 carboxyl-terminus. SnapGene and Clustal Omega were used to align UniProt-verified sequences for Hsp110 homologs from the following organisms: budding yeast (*S. cerevisiae*), fruit fly (*D. melanogaster*), human (*H. sapiens*), orangutan (*P. abelii*), bovine (*B. taurus*), Chinese hamster (*C. griseus*), mouse (*M. musculus*), and rat (*R. norvegieus*). Consensus sequence indicated in **bold**. Residues included in the consensus sequence are present in more than 50% of aligned species. Amino acids matching the consensus sequence are highlighted in pink. Bars indicate percent conservation at a specific amino acid position. red: 75-100%; orange: 50-75%; blue <50%.

**Figure S6:**
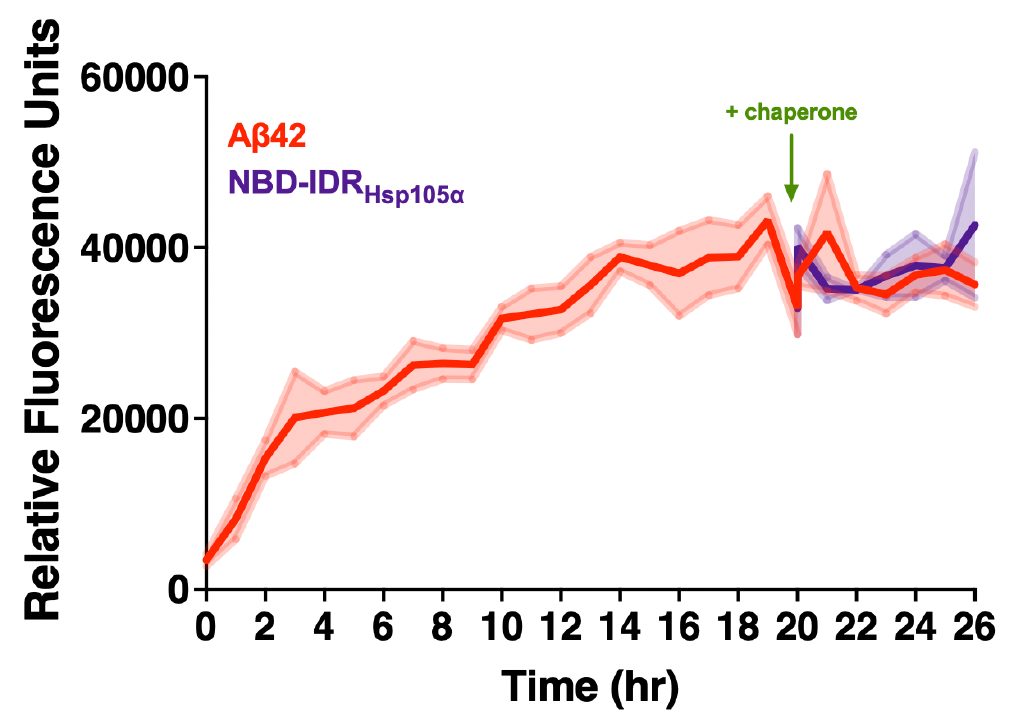
NBD-IDR_Hsp105α_ does not disassemble pre-formed Aβ42 fibrils. 2 μM Aβ42 was incubated alone for 20 hr as described in Materials and Methods, after which 4 μM NBD-IDR_Hsp105α_ was added to the reaction and allowed to incubate an additional 6 hr. Bolded lines are the average of three replicates for each condition while the shaded region represents standard error of the mean (SEM).

**Figure S7:**
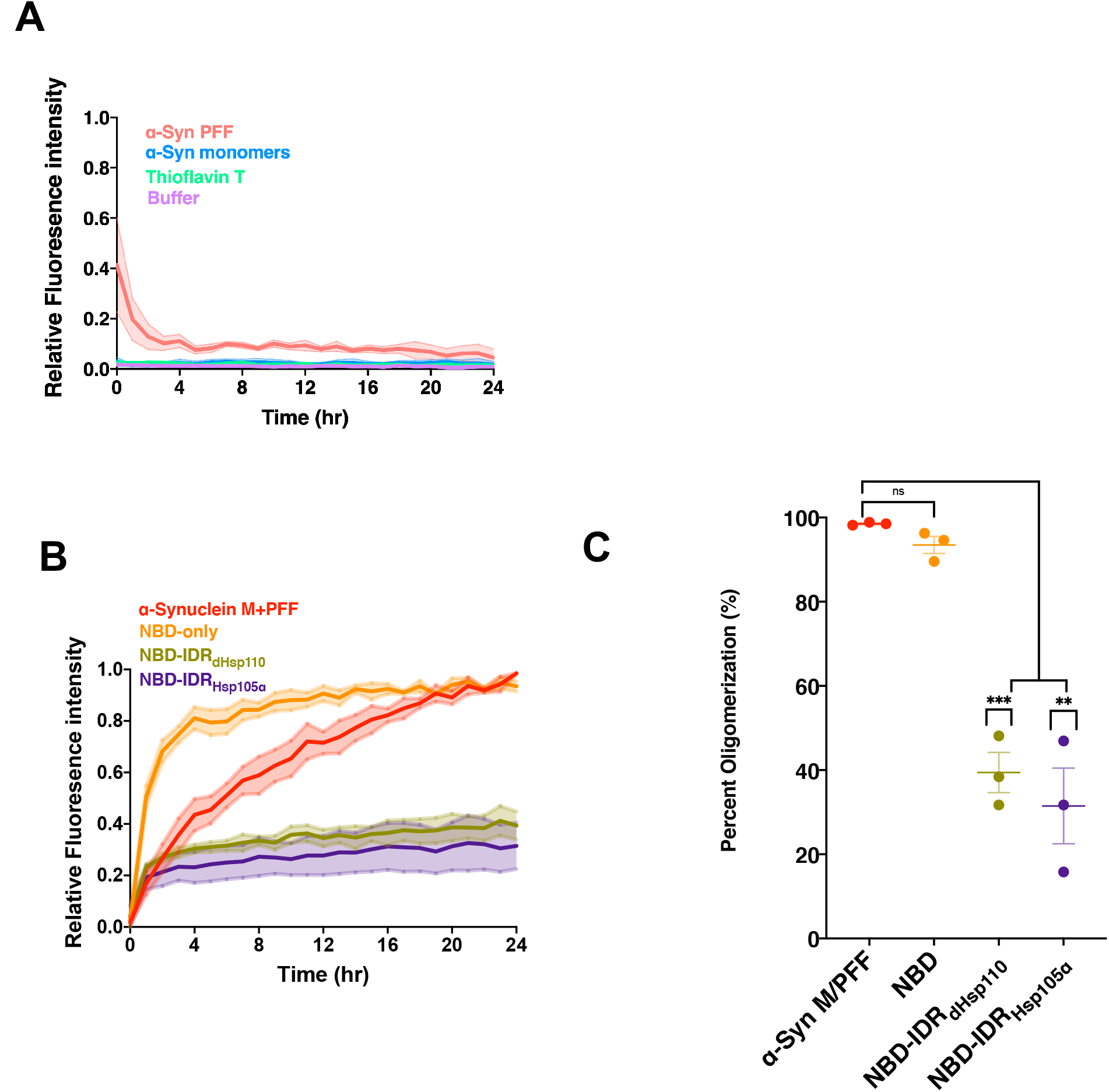
NBD-IDRs prevent α-synuclein oligomerization *in vitro*. **A**. Controls for thioflavin T binding assays and α-synuclein oligomerization. 2 μM α-synuclein monomers and 1 μM α-synuclein pre-formed fibrils were incubated individually in thioflavin T and Tris NaCl buffer. 5 μM thioflavin T was incubated alone in Tris NaCl buffer and fluorescence detected as described in Materials and Methods. Bolded lines are the average of three replicates for each condition while the shaded region represents standard error of the mean (SEM) **B**. 2 μM of α-synuclein monomers and 1 μM pre-formed fibrils incubated together (α-synuclein M+PFF) or with 4 μM of respective chaperone: NBD, NBD-IDR_dHsp110_, NBD-IDR_Hsp105α_ were incubated for 24 hr and thioflavin T fluorescence detected as described in Materials and Methods. Bolded lines are the average of three replicates for each condition while the shaded region represents standard error of the mean (SEM). **C**. End point measurements of each condition were taken from (B) and divided by the α-Synuclein M+PFF measurement within the respective replicate and converted to relative percentage. Group differences were analyzed using Welch’s *t*-test. *, p=0.05; **, p=0.005; ***, p=0.0005; ****, p=0.00005.

**Figure S8:**
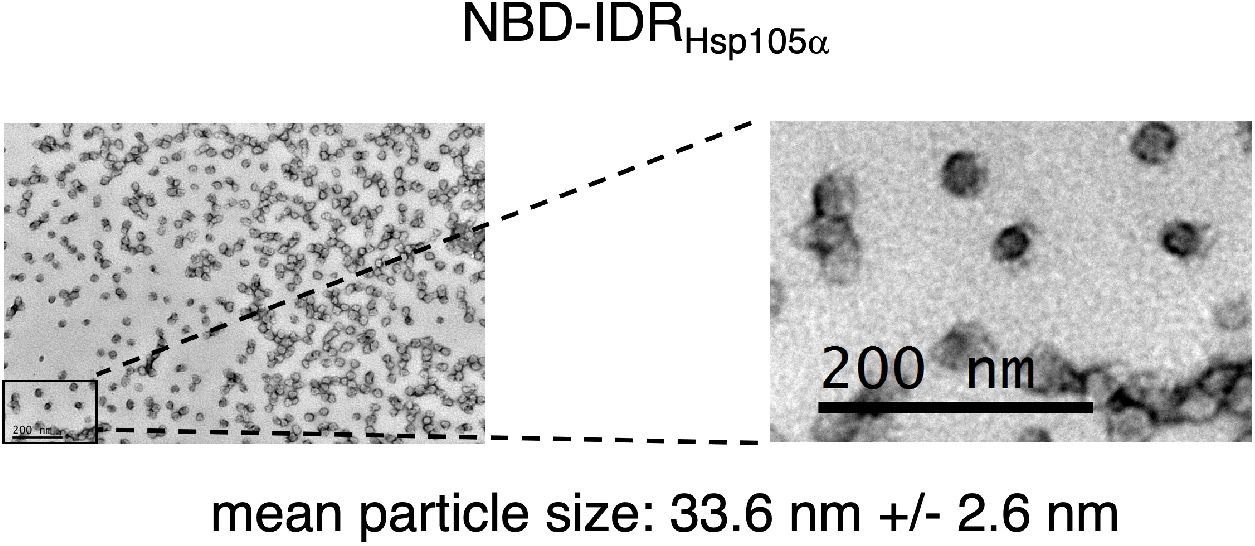
NBD-IDR_Hsp105α_ forms globular particles *in vitro*. 4 μM NBD-IDR_Hsp105α_ was incubated without substrate as described for Fig. 5 and imaged using transmission electron microscopy. Endpoint samples from the thioflavin T binding assay were recovered, negative stained and imaged via transmission electron microscopy, with or without additional chaperones as indicated. Particle size was measured (n=10) using ImageJ on the zoomed inset image. Scale bar= 200 nm.

## Notes

### Competing Interest Statement

The authors have declared no competing interest.

## References

1. Hartl FU, Bracher A, Hayer-Hartl M. 2011. Molecular chaperones in protein folding and proteostasis. Nature 475:324–332.

2. Lindquist S. 1986. The Heat-Shock Response. Annu Rev Biochem 55:1151–1191.

3. Morimoto RI. 1998. Regulation of the heat shock transcriptional response: cross talk between a family of heat shock factors, molecular chaperones, and negative regulators. Genes Dev 12:3788–3796.

4. Hipp MS, Park SH, Hartl UU. 2014. Proteostasis impairment in protein-misfolding and - aggregation diseases. Trends Cell Biol. Elsevier.

5. Hipp MS, Kasturi P, Hartl FU. 2019. The proteostasis network and its decline in ageing. Nat Rev Mol Cell Biol 20:421–435.

6. Labbadia J, Morimoto RI. 2015. The biology of proteostasis in aging and disease. Annu Rev Biochem 84:435–464.

7. Morimoto RI. 2008. Proteotoxic stress and inducible chaperone networks in neurodegenerative disease and aging. Genes Dev 22:1427–1438.

8. Gestwicki JE, Garza D. 2012. Protein quality control in neurodegenerative disease, p. 327–353. In Progress in Molecular Biology and Translational Science.

9. Kim YE, Hosp F, Frottin F, Ge H, Mann M, Hayer-Hartl M, Hartl FU. 2016. Soluble Oligomers of PolyQ-Expanded Huntingtin Target a Multiplicity of Key Cellular Factors. Mol Cell 63:951–964.

10. Knowles TPJ, Vendruscolo M, Dobson CM. 2014. The amyloid state and its association with protein misfolding diseases. Nat Rev Mol Cell Biol 15:384–396.

11. Olzscha H, Schermann SM, Woerner AC, Pinkert S, Hecht MH, Tartaglia GG, Vendruscolo M, Hayer-Hartl M, Hartl FU, Vabulas RM. 2011. Amyloid-like Aggregates Sequester Numerous Metastable Proteins with Essential Cellular Functions. Cell 144:67–78.

12. Taylor RC, Dillin A. 2011. Aging as an Event of Proteostasis Collapse. Cold Spring Harb Perspect Biol 3:1–17

13. Nagy M, Fenton WA, Li D, Furtak K, Horwich AL. 2016. Extended survival of misfolded G85R SOD1-linked ALS mice by transgenic expression of chaperone Hsp110. Proc Natl Acad Sci 113:5424–5428.

14. Wentink AS, Nillegoda NB, Feufel J, Ubartaitė G, Schneider CP, De Los Rios P, Hennig J, Barducci A, Bukau B. 2020. Molecular dissection of amyloid disaggregation by human HSP70. Nature 587:483–488.

15. Song Y, Nagy M, Ni W, Tyagi NK, Fenton WA, Lopez-Giraldez F, Overton JD, Horwich AL, Brady ST. 2013. Molecular chaperone Hsp110 rescues a vesicle transport defect produced by an ALS-associated mutant SOD1 protein in squid axoplasm. Proc Natl Acad Sci 110:5428–5433.

16. Kuo Y, Ren S, Lao U, Edgar BA, Wang T. 2013. Suppression of polyglutamine protein toxicity by co-expression of a heat-shock protein 40 and a heat-shock protein 110. Cell Death Dis 4:e833–e833.

17. Zhang S, Binari R, Zhou R, Perrimon N. 2010. A genomewide RNA interference screen for modifiers of aggregates formation by mutant huntingtin in drosophila. Genetics 184:1165–1179.

18. Taguchi Y V., Gorenberg EL, Nagy M, Thrasher D, Fenton WA, Volpicelli-Daley L, Horwich AL, Chandra SS. 2019. Hsp110 mitigates α-synuclein pathology in vivo. Proc Natl Acad Sci 116:24310–24316.

19. Eroglu B, Moskophidis D, Mivechi NF. 2010. Loss of Hsp110 Leads to Age-Dependent Tau Hyperphosphorylation and Early Accumulation of Insoluble Amyloid β. Mol Cell Biol 30:4626–4643.

20. Liu Q, Hendrickson WA. 2007. Insights into Hsp70 Chaperone Activity from a Crystal Structure of the Yeast Hsp110 Sse1. Cell 131:106–120.

21. Polier S, Hartl FU, Bracher A. 2010. Interaction of the Hsp110 Molecular Chaperones from S. cerevisiae with Substrate Protein. J Mol Biol 401:696–707.

22. Bracher A, Verghese J. 2015. The nucleotide exchange factors of Hsp70 molecular chaperones. Front Mol Biosci 2:10.

23. Dragovic Z, Broadley SA, Shomura Y, Bracher A, Hartl FU. 2006. Molecular chaperones of the Hsp110 family act as nucleotide exchange factors of Hsp70s. EMBO J 25:2519–2528.

24. Abrams JL, Verghese J, Gibney PA, Morano KA. 2014. Hierarchical Functional Specificity of Cytosolic Heat Shock Protein 70 (Hsp70) Nucleotide Exchange Factors in Yeast. J Biol Chem 289:13155–13167.

25. Garcia VM, Nillegoda NB, Bukau B, Morano KA. 2017. Substrate binding by the yeast Hsp110 nucleotide exchange factor and molecular chaperone Sse1 is not obligate for its biological activities. Mol Biol Cell 28:2066–2075.

26. Oh HJ, Chen X, Subjeck JR. 1997. hsp110 Protects Heat-denatured Proteins and Confers Cellular Thermoresistance. J Biol Chem 272:31636–31640.

27. Oh HJ, Easton D, Murawski M, Kaneko Y, Subjeck JR. 1999. The chaperoning activity of hsp110: Identification of functional domains by use of targeted deletions. J Biol Chem 274:15712–15718.

28. Xu X, Sarbeng EB, Vorvis C, Kumar DP, Zhou L, Liu Q. 2012. Unique peptide substrate binding properties of 110-kDa heat-shock protein (Hsp110) determine its distinct chaperone activity. J Biol Chem 287:5661–5672.

29. Raviol H, Sadlish H, Rodriguez F, Mayer MP, Bukau B. 2006. Chaperone network in the yeast cytosol: Hsp110 is revealed as an Hsp70 nucleotide exchange factor. EMBO J 25:2510–2518.

30. Garcia VM, Nillegoda NB, Bukau B, Morano KA. 2017. Substrate binding by the yeast Hsp110 nucleotide exchange factor and molecular chaperone Sse1 is not obligate for its biological activities. Mol Biol Cell 28:2066–2075.

31. McGurk L, Berson A, Bonini NM. 2015. Drosophila as an in vivo model for human neurodegenerative disease. Genetics 201:377–402.

32. Moloney A, Sattelle DB, Lomas DA, Crowther DC. 2010. Alzheimer’s disease: Insights from Drosophila melanogaster models. Trends Biochem Sci 35:228–235.

33. Sang TK, Jackson GR. 2005. Drosophila models of neurodegenerative disease. NeuroRx 2:438–446.

34. Garcia VM, Rowlett VW, Margolin W, Morano KA. 2016. Semi-automated microplate monitoring of protein polymerization and aggregation. Anal Biochem 508:9–11.

35. Shaner L, Trott A, Goeckeler JL, Brodsky JL, Morano KA. 2004. The function of the yeast molecular chaperone Sse1 is mechanistically distinct from the closely related Hsp70 family. J Biol Chem 279:21992–22001.

36. Jones DT, Cozzetto D. 2015. DISOPRED3: Precise disordered region predictions with annotated protein-binding activity. Bioinformatics 31:857–863.

37. Ward JJ, McGuffin LJ, Bryson K, Buxton BF, Jones DT. 2004. The DISOPRED server for the prediction of protein disorder. Bioinformatics 20:2138–2139.

38. Boczek EE, Alberti S. 2018. One domain fits all: Using disordered regions to sequester misfolded proteins. J Cell Biol 217:1173–1175.

39. Tsai C, Aslam K, Drendel HM, Asiago JM, Goode KM, Paul LN, Rochet J-C, Hazbun TR. 2015. Hsp31 Is a Stress Response Chaperone That Intervenes in the Protein Misfolding Process. J Biol Chem 290:24816–24834.

40. Cox D, Ecroyd H. 2017. The small heat shock proteins αB-crystallin (HSPB5) and Hsp27 (HSPB1) inhibit the intracellular aggregation of α-synuclein. Cell Stress Chaperones 22:589–600.

41. Raju M, Santhoshkumar P, Sharma KK. 2018. Cell-Penetrating Chaperone Peptide Prevents Protein Aggregation and Protects against Cell Apoptosis. Adv Biosyst 2:1700095.

42. Webster JM, Darling AL, Uversky VN, Blair LJ. 2019. Small heat shock proteins, big impact on protein aggregation in neurodegenerative disease. Front Pharmacol 10:1047.

43. Sievers F, Higgins DG. 2018. Clustal Omega for making accurate alignments of many protein sequences. Protein Sci 27:135–145.

44. Trivedi R, Nagarajaram HA. 2019. Amino acid substitution scoring matrices specific to intrinsically disordered regions in proteins. Sci Rep 9:1–12.

45. Shorter J. 2011. The mammalian disaggregase machinery: Hsp110 synergizes with Hsp70 and Hsp40 to catalyze protein disaggregation and reactivation in a cell-free system. PLoS One 6:e26319.

46. Scior A, Buntru A, Arnsburg K, Ast A, Iburg M, Juenemann K, Pigazzini ML, Mlody B, Puchkov D, Priller J, Wanker EE, Prigione A, Kirstein J. 2018. Complete suppression of Htt fibrilization and disaggregation of Htt fibrils by a trimeric chaperone complex. EMBO J 37:282–299.

47. Glenner GG, Wong CW. 1984. Alzheimer’s disease: Initial report of the purification and characterization of a novel cerebrovascular amyloid protein. Biochem Biophys Res Commun 120:885–890.

48. Kang J, Lemaire HG, Unterbeck A, Salbaum JM, Masters CL, Grzeschik KH, Multhaup G, Beyreuther K, Müller-Hill B. 1987. The precursor of Alzheimer’s disease amyloid A4 protein resembles a cell-surface receptor. Nature 325:733–736.

49. Selkoe DJ. 1991. The molecular pathology of Alzheimer’s disease. Neuron 6:487–498.

50. Bitan G, Kirkitadze MD, Lomakin A, Vollers SS, Benedek GB, Teplow DB. 2003. Amyloid-beta protein (Abeta) assembly?: Abeta 40 and Abeta 42 oligomerize through distinct pathways. Proc Natl Acad Sci U S A 100:330–335.

51. Yakubu UM, Catumbela CSG, Morales R, Morano KA. 2020. Understanding and exploiting interactions between cellular proteostasis pathways and infectious prion proteins for therapeutic benefit. Open Biol 10:200282.

52. Jarrett JT, Berger EP, Lansbury PT. 1993. The Carboxy Terminus of the β Amyloid Protein Is Critical for the Seeding of Amyloid Formation: Implications for the Pathogenesis of Alzheimer’s Disease. Biochemistry 32:4693–4697.

53. Gade Malmos K, Blancas-Mejia LM, Weber B, Buchner J, Ramirez-Alvarado M, Naiki H, Otzen D. 2017. ThT 101: a primer on the use of thioflavin T to investigate amyloid formation. Amyloid 24:1–16.

54. Xue C, Lin TY, Chang D, Guo Z. 2017. Thioflavin T as an amyloid dye: Fibril quantification, optimal concentration and effect on aggregation. R Soc Open Sci 4:160696.

55. Ota H, Fukuchi S. 2017. Sequence conservation of protein binding segments in intrinsically disordered regions. Biochem Biophys Res Commun 494:602–607.

56. Oldfield CJ, Dunker AK. 2014. Intrinsically disordered proteins and intrinsically disordered protein regions. Annu Rev Biochem 83:553–584.

57. Dyson HJ, Wright PE. 2005. Intrinsically unstructured proteins and their functions. Nat Rev Mol Cell Biol. 6: 197–208.

58. Uversky VN. 2019. Intrinsically disordered proteins and their “Mysterious” (meta)physics. Front Phys 7:8–23.

59. Haslbeck M, Vierling E. 2015. A first line of stress defense: Small heat shock proteins and their function in protein homeostasis. J Mol Biol 427:1537–1548.

60. Escusa-Toret S, Vonk WIM, Frydman J. 2013. Spatial sequestration of misfolded proteins by a dynamic chaperone pathway enhances cellular fitness during stress. Nat Cell Biol 15:1231–1243.

61. Garvey M, Ecroyd H, Ray NJ, Gerrard JA, Carver JA. 2017. Functional amyloid protection in the eye lens: Retention of α-crystallin molecular chaperone activity after modification into amyloid fibrils. Biomolecules 7:1–18.

62. Barrott JJ, Haystead TAJ. 2013. Hsp90, an unlikely ally in the war on cancer. FEBS J 1. 280:1381–1396.

63. Dewji NN, Do C. 1996. Heat shock factor-1 mediates the transcriptional activation of Alzheimer’s β-amyloid precursor protein gene in response to stress. Mol Brain Res 35:325–328.

64. Davis AK, Pratt WB, Lieberman AP, Osawa Y. 2020. Targeting Hsp70 facilitated protein quality control for treatment of polyglutamine diseases. Cell Mol Life Sci 77:977–996.

